# Constricted cell migration causes nuclear lamina damage, DNA breaks, and squeeze-out of repair factors

**DOI:** 10.1101/035626

**Authors:** Jerome Irianto, Charlotte R. Pfeifer, Yuntao Xia, Avathamsa Athirasala, Irena L. Ivanovska, Roger A. Greenberg, Dennis E. Discher

## Abstract

Genomic variation across cancers scales with tissue stiffness: meta-analyses show tumors in stiff tissues such as lung and bone exhibit up to 100-fold more variation than tumors in soft tissues such as marrow and brain. Here, nuclear lamina damage and DNA double-strand breaks (DSBs) result from invasive migration of cancer cells through stiff constrictions. DSBs increase with lamin-A knockdown and require micro-pores sufficiently small for lamins to impede migration. Blebs in the vast majority of post-migration nuclei are enriched in lamin-A but deficient in lamin-B and an age-associated form of lamin-A. Validation of DSBs by an electrophoretic comet assay calibrates against a cancer line having nuclease sites engineered in chromosome-1, and DSB-bound repair factors in nuclei pulled into constrictions show folded chromatin orients, extends, and concentrates without fragmentation. Mobile repair proteins simultaneously segregate away from pore-condensed chromatin. Global squeeze-out of repair factors and loss with lamin-A-dependent rupture explains why overexpression of repair factors cannot rescue DSBs in migration through stiff constrictions, ultimately favoring genomic variation.

## Introduction

Nuclear stiffness has long been speculated to limit a cell’s ability to migrate through more small, stiff pores, and our recent studies of cancer cells^5^ indeed showed that partial knockdown of lamin-A softened nuclei several-fold, enhanced constricted migration of cancer cells by several-fold, and also promoted more rapid invasion *in vivo* of nearby normal tissue. Similar studies of others likewise showed enhanced migration *in vitro*^3,19^. The nuclear lamina has likely evolved to protect the genome and enhance cell viability. Deep knockdown of lamin-A followed by constricted migration in our previous studies^5^ indeed resulted in enhanced cell death by mechanism(s) consistent with DNA damage.

DNA damage can also lead to genomic variation, with ionizing radiation being one long-accepted physical source of DNA damage causing genomic variation. Extensive sequencing of DNA from many bulk tumors prompted a meta-analysis of genomic variation in order to search for evidence in support of a novel hypothesis that invasive migration through stiff surroundings could damage the nucleus, including DNA. Stiff tissues such as lung, skin, and bone indeed show tumors with orders of magnitude greater genomic variation compared to soft tissues such as brain and marrow (**Fig. 1A**). Tissue stiffness depends in a power law fashion on the concentration of fibrous collagens^17^, and tissue stiffness is mechano-sensed by cells to cause many changes in cell phenotype, including a scaled increase in lamin-A levels that fortifies the nucleus against physical stress^17^. Quantitative analysis versus tissue stiffness shows both the extent of genomic variation across cancers and lamin-A levels are fit by similar power laws (exponents in the range of 0.6 – 1.0), and so we sought to quantify any damage to the nuclear lamina and DNA in relation to lamin-A levels after constricted migration of cancer cells.

**Figure 1.**
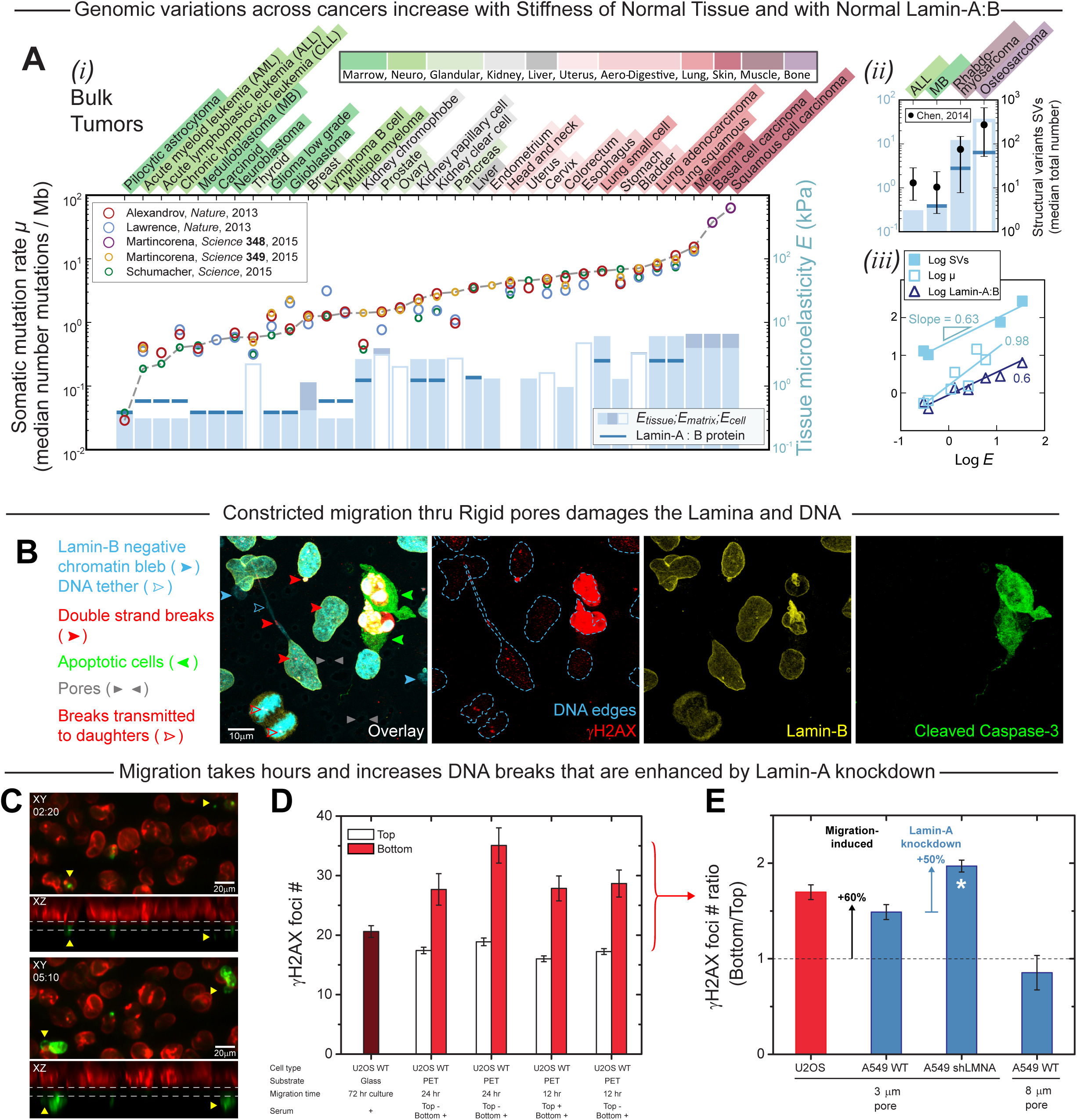
**(A)** Number of somatic mutations varies between different cancers^1–2, 8, 10–11, 15^. **(*i*)** Mutation rate correlates with tissue stiffness: softer tissues (green) such as marrow and brain seem to have ~100-fold fewer somatic mutations than the stiffer ones (red) like liver, lung or skin. While tissue stiffness scales with lamin-A:B ratio^17^. **(*ii*)** Similarly, structural variations also increase with tissue stiffness, highest in bone cancer. **(*iii*)** Structural variants, tissue microelasticity and lamin-A:B ratio scales with tissue stiffness. **(B)** A549 shLMNA still divide after constricted migrating through 3 μm Transwell membrane. Some cells also apoptose as shown by the cleaved caspase-3 and high level of γH2AX, a marker for DNA damage. Some level of γH2AX staining was also observed in the non-apoptosing cells, indicating a certain level of DNA damage. Some nuclei have parts of DNA that are not covered by the lamina: chromatin bleb and DNA tether. **(C)** Time-lapse confocal imaging of A549 shLMNA + GFP-Lamin-A^S22A^ (lamin-A with a non-phosphorylatable mutations at Serine 22) shows that it take approximately 3 hours for a nucleus to fully migrate thru the 3 μm constriction (hr:min, Red = nuclei on the top of the membrane, Green = nulcei passed through the constriction to the bottom of the membrane). Arrows showing the chromatin blebs observed in the fixed samples. **(D)** Migration through the 3 μm pores increased the number of γH2AX foci in U2OS cells, indicative of increased DSBs. Dark red bar shows the number of γH2AX foci in U2OS on conventional glass culture, PET = polyester 3 μm transwell membrane. Migration time and serum level do not significantly alter this migration induced DSB enhancement (n = 51–385 nuclei per condition). **(E)** Pore migration also induced DSB enhancement for A549s with normal lamin-A level (A549 WT), which is further increased by knocking down lamin-A level (A549 shLMNA). Enhancement of DSBs was not observed in cells migrated through bigger pores (8 μm) (N = 4-5, n = 51-578 per condition per experiment, mean±SEM, \**p*<0.001)

## Results and Discussion

In standard 2D cultures, basal levels of damage to the nuclear lamina and to DNA are expected to be low for cancer lines derived from relatively stiff tissues, including the two lines studied here: the A549 human-derived lung carcinoma line and the U2OS human osteosarcoma line. However, at the bottom of Transwell membranes with highly constrictive micro-pores (3 μm diameter pores)^5^, migration causes damage to both the lamina and to DNA (**Fig. 1B**). Blebs of chromatin that are not covered by lamin-B are very common for both cell types, and lamin-B-negative chromatin tethers are evident for A549 cancer cells. Double strand breaks (DSBs) marked by γH2AX foci are seen in nearly all nuclei, and apoptotic cells with the cleaved caspase-3 throughout the cell show very high γH2AX. Dividing cells positive for γH2AX staining also suggest DSBs are transmitted to daughter cells^9^. Time-lapse confocal imaging of A549 cells shows constricted migration across the filters takes ~3 hours (**Fig. 1C**), which is a significant timescale relative to repair of DSBs caused by ionizing radiation. Nuclear blebs with compromised lamina structure are also observed in cells expressing low levels of GFP-Lamin-A^S22A^, confirming results obtained from imaging of fixed samples.

Quantitation of γH2AX foci shows similar numbers on the Tops of Transwells as in conventional 2D cultures (**Fig. 1D**). However, γH2AX foci number in U2OS nuclei on the Transwell Bottoms are always significantly greater than on the Top, independent of migration time and serum gradient, and consistent with migration-induced DNA damage. A549 nuclei show a similar ~50–60% increase after constricted migration and also show lamin-A knockdown increases DNA damage (**Fig. 1E**), consistent with a protective role for lamin-A^5^. Importantly, DSBs do not increase for cells migrating through Transwells having large pores (8 μm) for which migration is independent of lamin-A^5^.

Blebbed nuclei post-migration show segregation of lamins (**Fig. 2A**) for the vast majority of both U2OS and A549 nuclei (**Fig. 2B**). Blebbing is independent of lamin-A but requires small pores rather than 8 μm pores. Blebs are enriched in lamin-A, which is in a dilated meshwork as simulated recently^4^, but blebs are deficient in lamin-B and DNA (**Fig. 2C**). The observations are consistent with blebs observed recently within a small minority of cells expressing the aging mutant of lamin-A, Progerin, in 2D cultures^18^. Unlike lamin-A, both Progerin and lamin-B are farnesylated and immobile on the nuclear envelope. Lamin-A knockdown doubles bleb numbers on the Transwell Tops and increases the frequency of chromatin tethers (of length >10 μm) before and after constricted migration (**Fig. 2D**), but such tethers are only evident for A549 cells, which indicates that DNA damage to U2OS cells (Fig. 1D,E) is independent of any tethering process. Ectopic expression of wild-type GFP-lamin-A leads to the same lamin-A localization to nuclear blebs post-migration (**Fig. 2E**), whereas GFP-Progerin does not enrich, behaving similar to lamin-B. Aging is of course the main risk factor for human cancer, and progeria cells are well known to exhibit abnormally high DNA damage.

**Figure 2.**
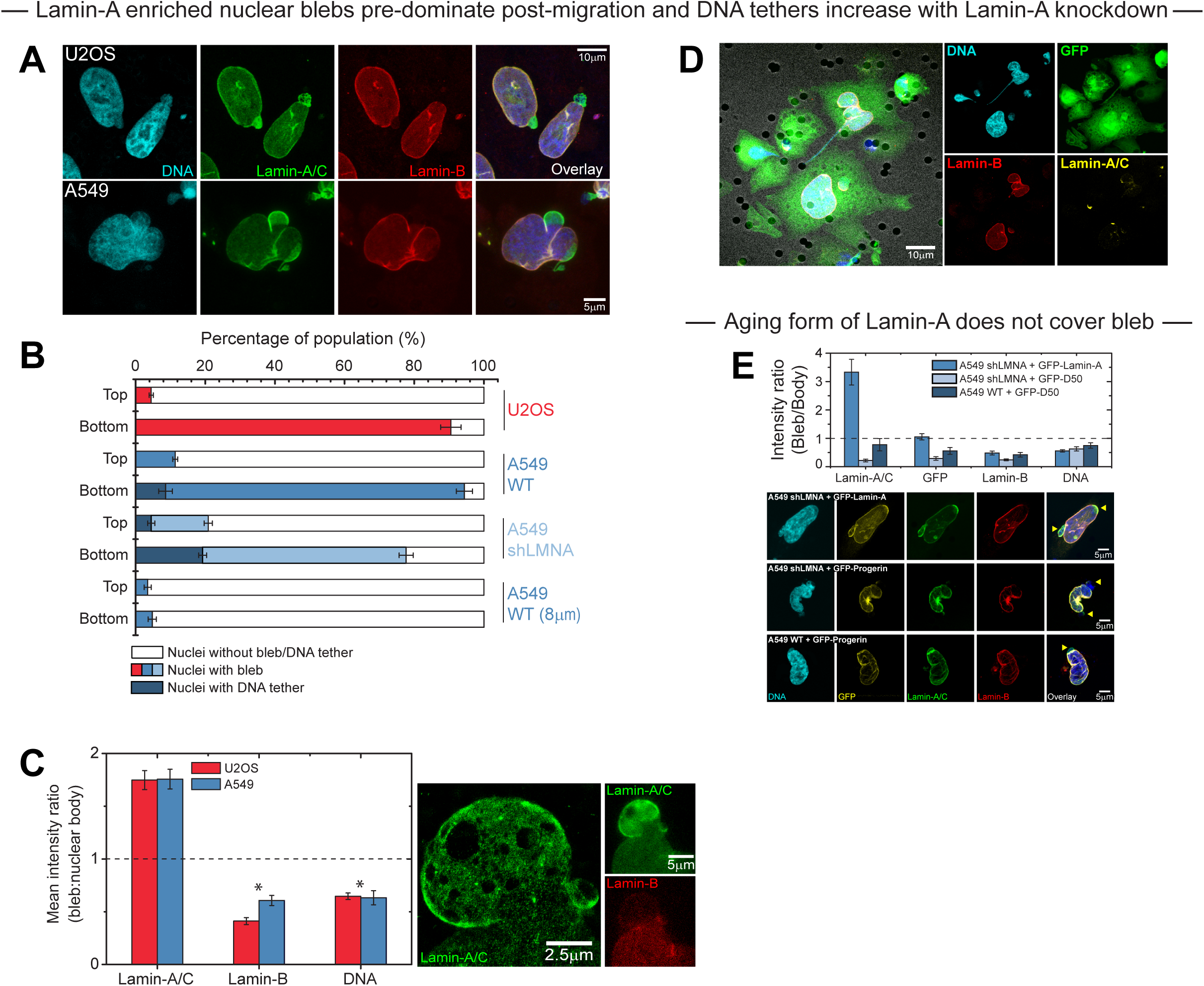
**(A)** Nuclear blebs formation in wild-type A549 and U2OS cells was observed after migration through 3 μm pores, **(B)** which dominate the population of migrated cells (>90%), not observed for the bigger pores. Increased number of DNA tether was also observed with A549s after migration, which seems to increase with lamin-A knockdown. **(C)** Nuclear blebs are clearly enriched in lamin-A and deficient in both lamin-B and DNA (n = 44-57 nuclei). Super resolution image of the nuclear bleb, showing dilated mesh network of Lamin-A,C (taken with Leica SR GSD). **(D)** The observed DNA tethers always lead to a pore and still covered by the cytoplasm, as shown by the GFP signal. **(E)** Enriched lamin-A blebs are rescued by GFP-Lamin-A over-expression, while progerin (GFP-Progerin) behave like the deficient lamin-B.

In order to validate the DNA damage results above and perhaps elucidate aspects of mechanisms, the U2OS cells used throughout this study were engineered^16^ with inducible and visualizable DNA damage in a p-arm locus inserted within one copy of Chromosome-1 (**Fig. 3A**). Even in 2D culture, induction of the mCherry-tagged nuclease produces correlations (*i*) between γH2AX intensity at the locus and nuclease intensity, and also *(ii)* between the projected areas of the DNA damage response factor GFP-53BP1and γH2AX at the locus. In an electrophoretic comet assay, the well-controlled cleavage of DNA by nuclease produces a large and expected shift in the DNA centroid toward the cathode in the majority of nuclei from 2D cultures (**Fig. 3B**). Importantly, both the proportion of nuclei with a centroid shift (above a threshold) and the mean shift in centroid are higher for nuclei taken from the Transwell Bottom relative to the Top of the same Transwell. This independent method of detecting DNA breaks confirms the post-migration increase in DNA damage based on γH2AX immunostaining (Fig. 1B,D,E) and also indicates a linear correlation between the results from the two methods (**Fig. 3C**).

**Figure 3.**
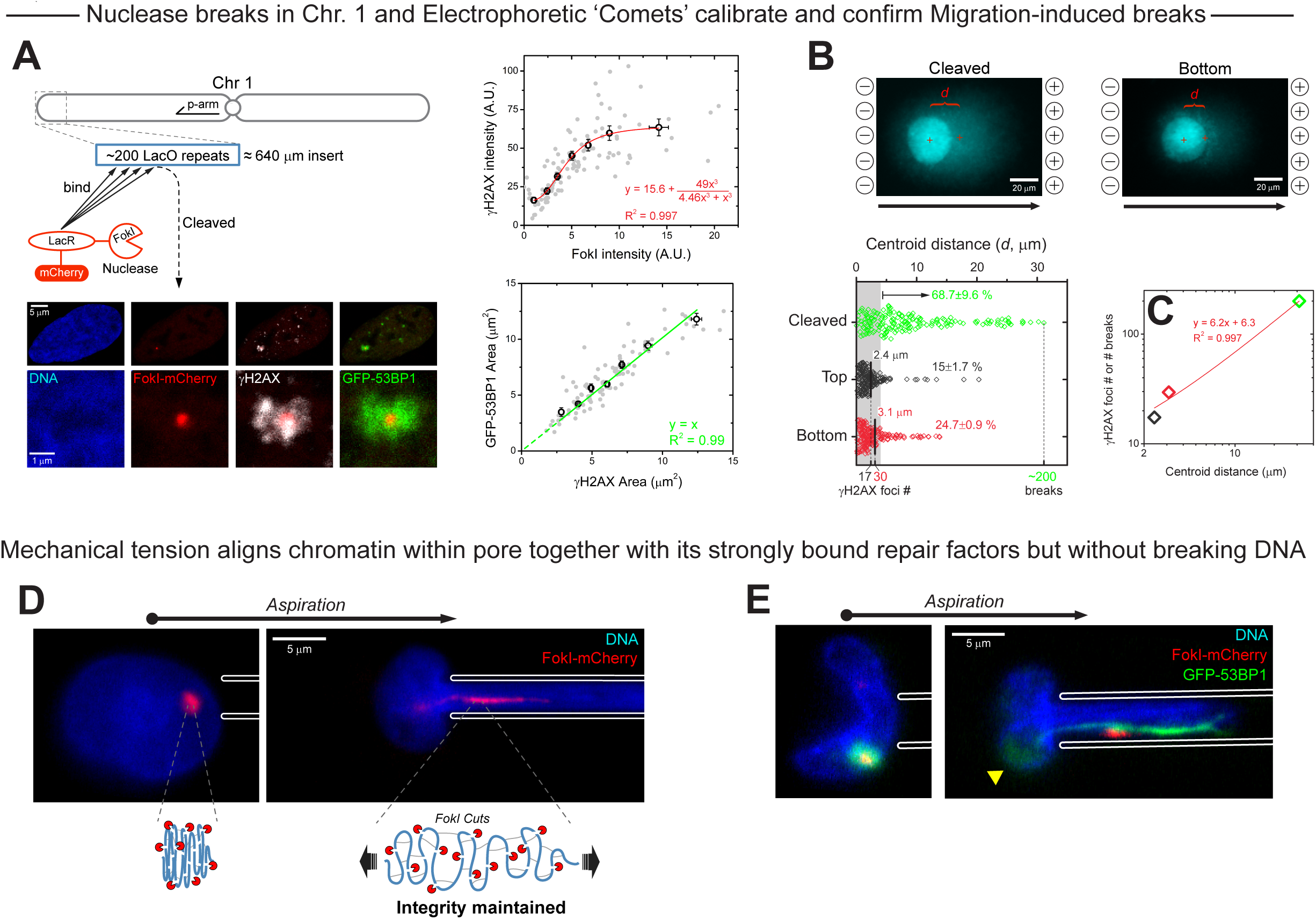
**(A)** U2OS with integrated lac operator transgene at the p-end of chromosome 1, DSBs can be induced by the expression of the integrated FokI-Lac Repressor-mCherry construct. FokI intensity correlates with γH2AX, while area of GFP-53BP1 at the damaged site is linearly correlated to γH2AX foci area. **(B)** Comet assay is sensitive enough to detect the induced DSBs at the inserted lac operator repeats (Green). The proportion of nuclei with centroid distance above the threshold is higher for the migrated samples (n = 176-199 nuclei per group, 3 experiments), confirming the increased DNA damage after migration shown by γH2AX staining in Fig. 2D,E. **(C)** The level of DNA damage was estimated from the centroid distance by using the earlier γH2AX data, revealing a linear correlation between the two. Allowing the estimation of DNA damage from the centroid distance. **(D)** Prior to micro-pipette aspiration, FokI contract was activated and the U2OS cells were treated with latrunculin-A. Aspiration with 3 μm pipette revealed that the damage site stretches and aligns as it goes into the constriction. **(E)** The repair scaffold as shown by GFP-53BP1 focus also behave the same, it stretches and aligns with the constriction. Some residual GFP-53BP1 was observed outside of the pipette (arrow).

Visualization of a chromatin break locus being pulled dynamically into a constricting pore was achieved by inducing the fluorescent nuclease and subsequently aspirating a suspended cell into a micropipette of ~3 μm diameter. The damage locus stretches and aligns as it enters the constriction (**Fig. 3D**), and extension is maximized to ~10 μm by both inhibiting the recruitment of DNA repair factors (with ATM kinase inhibitor, KU55933) and by de-condensing heterochromatin via a hypotonic pre-treatment^7^. Locus integrity is nonetheless maintained, suggesting chromatin cohesion factors maintain a highly folded chromatin structure. The ~10 μm extension is indeed only ~1% of the 640 μm locus length. The repair scaffold factor GFP-53BP1 (which depends on ATM kinase) which surrounds the locus (Fig. 3A, images) also stretches within the aligning constriction and mostly maintains continuity along a much longer length of the chromosome (**Fig. 3E**). However, a small pool of diffuse GFP-53BP1 is also evident outside of the micropipette, suggesting a segregation of unbound repair factor.

In the absence of nuclease induction, GFP-53BP1 is seen to be diffuse, consistent with nucleoplasm mobility, but micropipette aspiration largely segregates the GFP-53BP1 away from the chromatin pulled into the constriction (**Fig. 4A**). Chromatin condensation near the entrance of the constriction is indeed evident regardless of whether the nuclease was induced (with or without ATM inhibition Fig. 3D,E) or not (Fig. 4A). Some nuclei at large extension into the micropipette also show a nuclear bleb or rupture that is predictably rich in segregated GFP-53BP1. Chromatin bound proteins such as H2B and HP1α do not segregate and respond like DNA as quantified through an Intensity Ratio for GFP-protein just Inside/Outside the micropipette (**Fig. 4B**). Such an association with DNA agrees very well with observations above for GFP-53BP1 bound around the cleaved locus after nuclease induction (Fig. 3E). Micropipettes with large diameters do not show significant segregation of GFP-53BP1. Mechanistically, higher DNA intensity at the micropipette’s entrance indicates a condensation of chromatin caused by the constriction geometry, and such chromatin condensation will tend to sterically exclude nucleoplasmic factors and physically squeeze out all mobile proteins.

**Figure 4.**
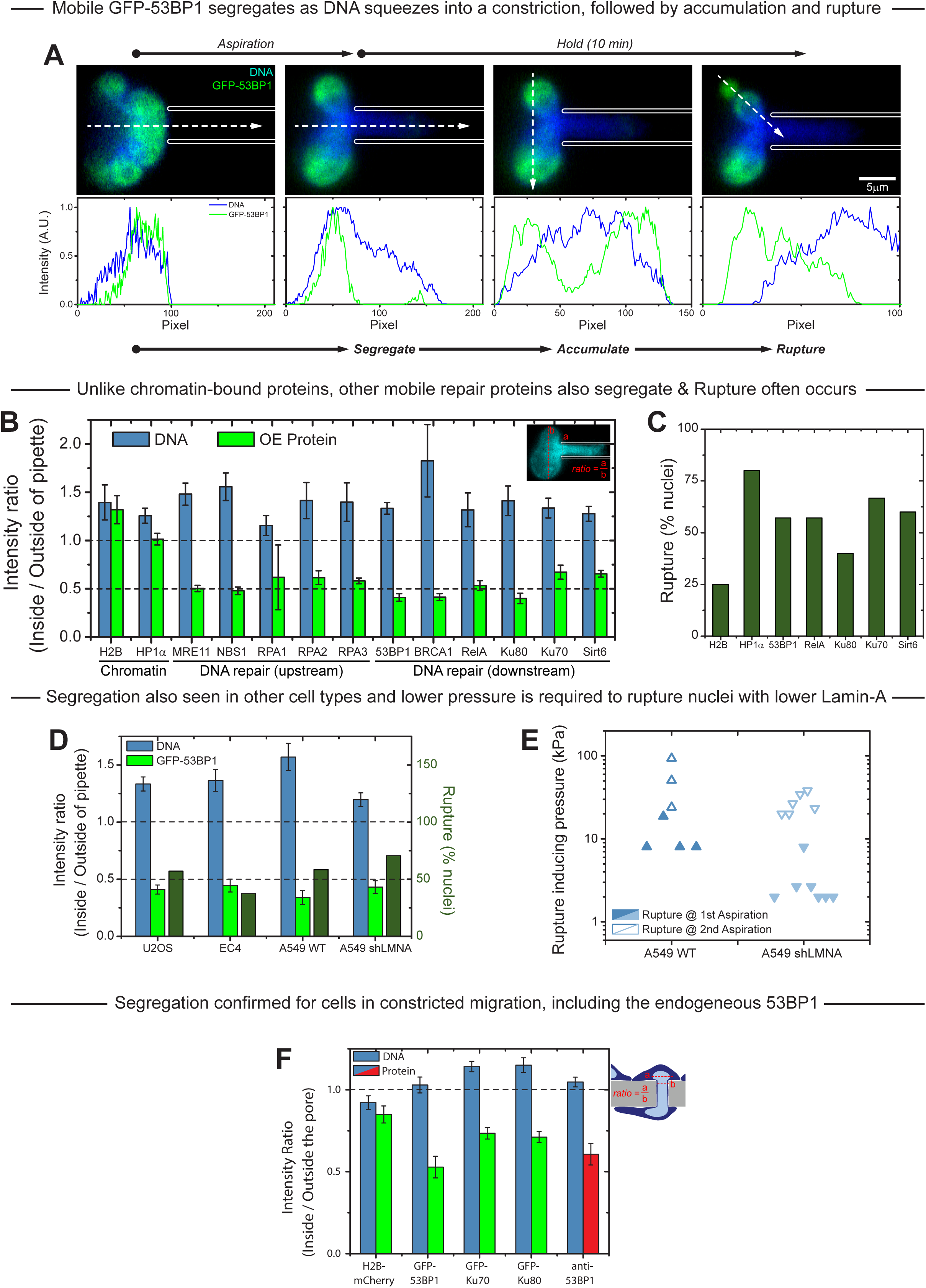
**(A)** Micropipette aspiration (~Ø3 μm) of intact U2OS, treated with latrunculin A. As the nucleus enter the pipette, mobile GFP-53BP1 segregates and accumulates outside the pipette, followed by rupture. **(B)** Segregation was observed in other mobile proteins, as shown by the intensity ratio. Higher DNA intensity at the pipette’s neck suggest a compaction caused by the constriction geometry, not allowing the mobile protein to go through. **(C)** Nuclear rupture was often observed. **(D)** Segregation and rupture of GFP-53BP1 with aspiration was also observed in other cell types: A549 and EC4, a liver cancer cell line. **(E)** Although the proportion of cells that rupture is similar for A549 WT and A549 shLMNA, lower pressure is required to induce the rupture in the lamin-A knockdowned cells. **(F)** Segregation of mobile protein was also observed in our Transwell migration samples as the nucleus enter the pore, which was also confirmed by the endogenous 53BP1.

All other mobile proteins indeed segregate similar to GFP-53BP1, based once again on the Inside/Outside intensity ratio (Fig. 4B). These include both upstream DNA damage response proteins like MRE11 and RPA^6^, and downstream factors like BRCA1 and Ku70 and Ku80. The dozen mobile proteins studied also include a transcription factor (RelA), an epigenetic factor (Sirt6), and the gene editing enzyme Cas9.

Rupture of the nuclear envelope was often observed (**Fig. 4C**), with histone-H2B showing the lowest probability of being lost in a rupture event – consistent with strong binding to chromatin. A physically unavoidable steric exclusion mechanism should and does also apply to GFP-53BP1 segregation and loss in rupture upon aspiration of other cancer cell types (**Fig. 4D**), including a mouse liver cancer line, EC4, and A549 cells +/-lamin-A knockdown. The latter results hint at more rupture events for low lamin-A, which prompted a more careful study of rupture as a function of aspiration pressure. Lower pressure is indeed required to cause rupture in the lamin-A knockdown cells (**Fig. 4E**), which is again indicative of a protective role for lamin-A even on the minute time scale of aspiration. Such a global loss of repair factors could explain why lamin-A knockdown cells show more DNA damage after migration through a Transwell with small and stiff micro-pores (Fig. 1E). Consistent with a close correlation between single cell aspiration studies and constricted migration, segregation of 53BP1 relative to DNA was also measured after Transwell migration as the nucleus either enters micro-pore or leaves it, both for ectopic GFP-53BP1 and immunostained, endogenous 53BP1 (**Fig. 4F**).

Confocal imaging of nuclei that are coming out of micro-pores show (in side-view) γH2AX foci throughout the nucleus (**Fig. 5A**), including near the lamina, but many DNA damage foci lack 53BP1, as confirmed by a count of foci (**Fig. 5A,B**). 53BP1 might thus be a limiting factor in DNA repair. GFP-53BP1 was therefore over-expressed prior to constricted migration (**Fig. 5C**). Despite the large excess of this DNA damage response protein, migration-induced DNA damage is not rescued (**Fig. 5D**). Overexpression of another DNA repair protein, Ku70, also does not rescue the DNA breaks (**Fig. 5E**).

**Figure 5.**
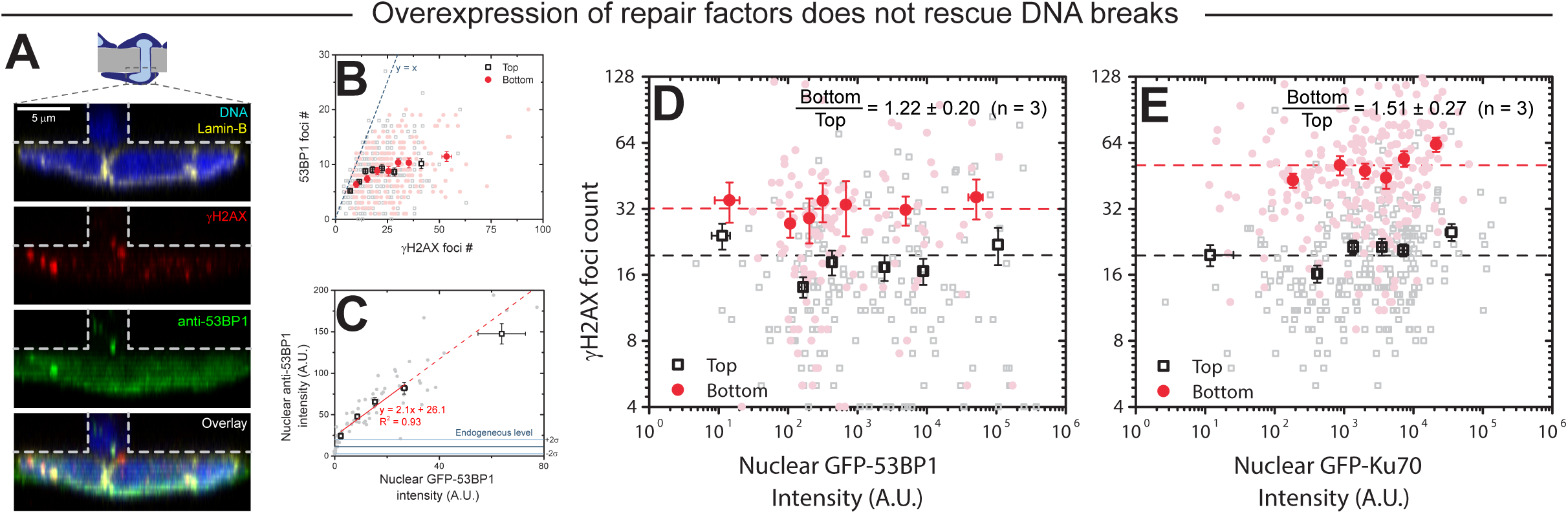
**(A)** Representative images of confocal projections at the XZ axis (side-view), showing that not all DNA damage sites covered by γH2AX are covered by 53BP1, as confirmed by **(B)** a count of foci. **(C)** GFP-53BP1 was over-expressed prior to Transwell migration. **(D, E)** Overexpression of neither 53BP1 nor Ku70, a DNA repair protein, can rescue migration-induced DNA damage.

The DNA damage response and repair pathway is certainly a multi-factorial pathway^13^, but segregation of all mobile factors during constricted migration could ultimately delay DNA repair. Accumulation of DNA damage and its mis-repair due to constricted migration through stiff micro-pores could thus contribute to genomic variation that our initial meta-analyses shows is much greater in stiffer tissues.

## Materials and methods

### Cell culture

Unless stated, A549, human lung cancer cell line, and U2OS, bone cancer cell line, were cultured in Ham’s F12 nutrient mixture and DMEM high glucose media (Gibco, Life Technologies), respectively, supplemented with 10% FBS and 1% penicillin/streptomycin (Sigma-Aldrich). Partial knock-down of Lamin-A in A549 cells were achieved by using shLMNA as described previously^5^. 256 lac operator repeats were integrated to the p-end of chromosome 1 of U2OS cells used in this study. DSBs on this site can be induced by the activation of endonuclease FokI-lac repressor-mCherry construct with the addition of 4-hydroxytamoxifen (Sigma) and Shield1 ligand (Clontech), as described previously^16^. When overexpression of a protein is involved, cells were transfected by Lipofectamine 2000 (Invitrogen, Life Technologies) for 24 hours prior to further experimentation.

### Transwell migration

Unless stated, for Transwell migration (Corning), cells were seeded at 300,000 cells/cm^2^ onto the top side of the membrane and left to migrate in normal culture condition for 24 hours. When immunostaining is involved, the membrane then fixed in 4% formaldehyde (Sigma) for 15 minutes, followed by permeabilization by 0.25% Triton-X (Sigma) for 10 minutes, blocked by 5% BSA (Sigma) and overnight incubation in various primary antibodies: lamin-A/C (Santa Cruz and Cell Signalling), Lamin-B (Santa Cruz), cleaved caspase-3 (Abcam), γH2AX (Milipore) and 53BP1 (Abcam). Finally, the primary antibodies were tagged with the corresponding secondary antibodies for 1.5 hours. When mounting is involved, Prolong Gold antifade reagent was used (Invitrogen, Life Technologies). Confocal imaging was done in Leica TCS SP8 system, by either 63x/1.4 oil-immersion or 40x/1.2 water-immersion objectives. Commet assays of the migrated cells were carried out per manufacturer’s instruction (Cell Biolabs). Cells were detached from the Transwell by using 0.05% Trypsin-EDTA (Gibco, Life Technologies).

### Micropipette aspiration

Prior to aspiration experiments, cells were detached with Trypsin-EDTA and treated with 0.2 μg/mL latrunculin-A (Sigma) for 1 hour at 37^o^C as described previously^12^, and re-suspended in aspiration buffer (PBS with 1% BSA). Nuclei were stained with 8 μM Hoechst 33342 for 15 minutes. Epifluorexcence imaging was done with Nikon TE300, with a digital CCD camera (Roper Scientifice) and a 60x/1.25 oil-immersion objective. Various image quantification and processing were done with ImageJ^14^.

## Acknowledgment

The authors would like to thank Jiri Lukas (University of Copenhagen, Denmark) and Marc S. Wold (University of Iowa, IA) for various DNA damage response protein plasmids used in this study. The authors in this study were supported by the National Cancer Institute of the National Institutes of Health under Award Number U54. The content is solely the responsibility of the authors and does not necessarily represent the official views of the National Institutes of Health

